# Characterization of Differences of Seed Endophytic Microbiome in Conventional and Organic Rice by Amplicon-based Sequencing and Culturing Methods

**DOI:** 10.1101/2023.10.10.561698

**Authors:** Sabin Khanal, Muhammad Imran, Xin-Gen Zhou, Sanjay Antony-Babu

## Abstract

The seed serves as the primary source of microbial inoculum for plant microbiota, playing a crucial role in establishing microbial populations in plants across subsequent generations, ultimately impacting plant growth and its overall health. Cropping conditions, especially farming practices, can influence the composition and functionality of the seed microbiome. Very little is known about the differences in seed microbiome between organic and conventional production systems. In this study, we characterized the endophytic microbial populations in seeds of rice grown under organic and conventional management practices through culture-dependent and independent analyses. The V4 region of 16S rRNA was used for bacterial taxa identification, and the ITS1 region was used in the identification of fungal taxa. Our results revealed significantly higher Shannon and Simpson indices for bacterial diversity in the conventional farming system whereas the fungal diversity was higher for observed, Shannon, and Simpson indices in the organic farming system. The cultivable endophytic bacteria were isolated and identified by the full-length 16S rRNA gene. There was no difference in culturable endophytic bacterial isolates in rice seeds grown under both conventional and organic farming systems. Among 33 unique isolates tested *in vitro*, three bacteria *Bacillus* sp. ST24, *Burkholderia* sp. OR5, and *Pantoea* sp. ST25, showed antagonistic activities against *Marasmius graminum, Rhizoctonia solani* AG4, and *R. solani* AG11, the fungal pathogens causing rice seedling blight.

**IMPORTANCE:** In this paper, we studied the differences in the endophytic microbial composition of rice seeds grown in conventional and organic farming systems. Our results demonstrate a greater bacterial diversity in conventional farming, while organic farming showcases a higher fungal diversity. Additionally, our research reveals the ability of seed bacterial endophytes to inhibit the growth of three fungal pathogens responsible for causing seedling blight in rice. This study provides valuable insights into the potential use of beneficial seed microbial endophytes for developing a novel microbiome-based strategy in the management rice diseases. Such an approach has the potential to enhance overall plant health and improve crop productivity.

---

Plant-associated microbial communities affect various plant traits (Shade et al. 2017). The plant microbiota can be acquired through either vertical transmission from the parent or horizontal transmission form the environment (Gundel et al. 2011). While the seed microbiota performs as a primary influencer for the overall plant microbiome (and hence the plant health), the seed microbiota can also play a direct role in seed germination and inducing tolerance to biotic and abiotic stresses (Goggin et al. 2015; Eyre et al. 2019). Multiple contributing factors can affect plant microbial community structures, such as environmental perturbation, soil composition and microbes, plant genotype, etc. (Hacquard 2016; Hamid et al. 2017). Among the environmental factors, farming practices play an important role in microbial community diversity (Reinhold-Hurek and Hurek 2011). The seed microbiomes are estimated to have more than 9,000 different microbial taxa, which can engage in synergistic, commensal, and potentially pathogenic interactions with the plant (Berg and Raajimakers 2018). Bacterial endophytes are among the most common microbes, with *Actinobacteria, Bacteroidetes, Firmicutes,* and *Proteobacteria* being the representative phyla (Barret et al. 2015; Bulgarelli et al. 2013; Links et al. 2014; Liu et al. 2012). Similarly, several fungal classes such as *Dothideomycetes, Eurotiomycetes, Leotiomycetes, Sardariomycetes,* and *Tremellomycetes* are also reported as endophytic fungal populations in seeds (Li et al. 2019). In the study presented here, we investigated seed microbiota of rice. Previous studies examining the influence of conventional and organic farming systems have primarily focused on microbial communities in the rhizosphere soil (Hartmann et al. 2015; Lupatini et al. 2017; Mader et al. 2002; Peltoneimi et al. 2021). These studies provide evidence of an augmented species number and enhanced diversity in the rhizosphere of organic production systems. Some studies have also shown the ability of seed microbial populations to protect seedlings from various soilborne pathogens (Beckstead et al. 2007; McArt et al. 2014). Seed endophytes with anti-fungal properties hold promise as potential to serve as effective biocontrol agents in agricultural settings (Truyens et al. 2014; Li et al. 2019).

Demand for organic rice has led to an almost six-fold increase in organic rice production in the US since 1995. Texas and California, together, account for 76% of this increase and stand as the largest organic rice-producing states. With increased production, it is critical to design, effective management strategies against crop loss due to disease, insects, and weeds. However, such management tools are currently unavailable for rice growers (Zhou et al. 2021a). The identification and development of effective biocontrol agents can offer rice farmers with such a tool for disease management, especially considering that the use of synthetic fungicides is prohibited for use in organic production systems. In addition, conventional rice growers are interested in adopting integrated disease management practices for rice production. Integrated use of biocontrol agents with conventional synthetic products can improve control efficacy, increase the longevity of the chemicals, and decrease the chemical load into the environment (Prajapati et al. 2020; Zhou et al. 2021b). Several bacterial taxa are known to have antagonistic properties against fungal pathogens and have the potential as disease biocontrol agents (Fira et al. 2018; Walterson and Stavrinides 2015). Many bacterial taxa are known to possess biocontrol properties against fungal pathogens of rice and other crops, which is especially true for members of the genera *Bacillus*, *Pantoea*, *Pseudomonas*, and *Streptomyces* (Kumar et al. 2011; Zhou et al. 2021b). These bacterial species target various fungal pathogens ranging from soil borne pathogens to foliar pathogens (Bardin et al. 2003; Hinarejos et al. 2016; Hu et al. 2007; Huang et al. 2012; Luo et al. 2015; Meena and Kanwar 2015; Waewthongrak et al. 2015; Zhoa et al. 2014).

Seedling blight is an important disease in rice in the southern US. It causes irregular, thin stand and weakened plants, resulting in significant stand loss and yield reduction (Zhou and Jo 2023). Dry seedling is the predominant method for rice cultivation in Texas and other southern states. Early plantings are widely practiced in Texas and Louisiana, allowing to produce a second crop (ratoon) while reducing the likelihood of heat stress and late-season disease occurrence. In such production conditions, seedling blight has emerged as the primary factor contributing to stand loss. This is primarily due to the onset of cold soil temperatures during early stages, which facilitates the development of seedling blight (Gaire et al. 2022; Rush 2018). Seedling blight causes pre- and post-emergence damping-off of rice plants. Although some blighted seedlings survive, they still suffer from poor vigor and have stunted growth. Based on the most recent seedling disease surveys, *Rhizoctonia solani* AG11 is the most abundant fungal pathogen that causes seedling blight in the southern US (Gaire et al. 2022). *R. solani* AG4 and *Marasminus graminum,* the two new fungal pathogens, also causing the seedling blight (Gaire et al. 2020, 2021, 2023). Currently, farmers heavily rely on fungicide seed treatment for the control of these seedling diseases (Jones and Carling 2007; Zhou and Jo 2023).

The objective of this study is to examine whether the farming practices, conventional and organic farming, affects seed microbial composition, and to what could be the potential major microbial players. Our goal is to build a foundational study that can lead to larger studies across different geographical regions across the world. To the best of our knowledge, this is the first study examining the diversity of seed endophytic microbial communities in the spermosphere of conventionally and organically grown rice. In addition, we also aimed to identify naturally inhabiting microorganisms that held potential for managing seedling blight in rice. Our results indicate that the diversity of endophytic bacterial communities in rice seeds was significantly higher in the conventional farming system compared to the organic system. Conversely, the diversity of endophytic fungal communities was greater in organically grown rice seeds. Furthermore, we observed that seed endophytic bacteria exhibited antagonistic properties against three seedling blight-causing pathogens in rice: *M. graminum, R. solani* AG11, and *R*. *solani* AG4.

## MATERIALS AND METHODS

### Collection of seed samples

In this study, rice seed samples were all collected in 2020 from two rice fields in half kilometer apart at the Texas A&M AgriLife Research Center, Beaumont, Texas. One field was under conventional management, while the other followed organic practices. The conventionally managed field’s coordinates were 30°03’37.9” N 94°17’34.7” W, while the organically managed field’s coordinates were 30°03’55.2” N 94°17’51” W. Both fields shared the same type of soil, classified as league-type, with the following composition 3% sand, 32% silt, 64% clay, 4% organic matter, and pH 5.5. The conventional field had been cropped to conventional rice every other year for more than 20 years. A typical growing season for the conventional field consisted of one or two applications of fungicides (azoxystrobin, a QoI complex 3 respiration inhibitor fungicide, 6.9 liter/ha and/or propiconazole, a sterol biosynthesis inhibitor fungicide, 7.3 liter/ha), one application of an insecticides (Zeta-cypermethrin, a group 3A insecticide, 2.9 liter/ha), and three applications of herbicides (clomazone, 0.9 liter/ha; halosulfuron, 0.09 liter/ha; and penoxsulam, 2.0 liter/ha). Synthetic fertilizers, mostly urea (46-0-0, N-P-K), were applied at 202 kg/ha to the field in each of cropping seasons. On the other hand, the organic field followed organic management practices and had been certified for organic rice production since 2007. In this organic approach, no synthetic chemicals such as fungicides, insecticides, herbicides or fertilizers (urea) were used. Instead, the organic-certified soil amendments Nature Safe (13-0-0, N-P-K), Rhizogen (7-2-1), and AgriRecycle (4-2-3) were utilized, along with winter cover crops such as white clover, purple clover or annual ryegrass to provide nutrients for the organic rice crops. For the 2020 cropping season, the study involved both conventional and organic trials, each trial was conducted using a randomized complete block design with four replications. The plots were4.9 m long and 1.3 m wide, consisting of eight rows with a spacing of 18 cm. Both field trials were drill seeded with the same rice cultivar Presidio at 134 kg/ha on April 27^th^. However, the seed used for the organic rice trial was harvested from the previous year’s organic crop. Both trials were managed following local conventional or organic rice production recommendations on fertility, irrigation, pest control, and other agronomical practices (Zhou et al. 2021a; Zhou and Jo 2023). Briefly, for the conventional trial, plots received 56 kg N/ha of the fertilizer urea on May 15^th^, and 168 kg N/ha of the fertilizer on June 7^th^. Permanent flood was established on June 7^th^. Plots were treated with the herbicides Command (clomazone, 0.94 liter/ha) and Halomax (halosulfuron, 0.09 liter/ha) on April 25^th^ for weed control. However, for the organic trial, plots received 56 kg N/ha of organic-certified fertilizer Nature Safe on May 14^th^, 168 kg N/ha on June 1^st^, and 168 kg N/ha on July 1^st^. Permanent flood was established on June 1^st^. Both trials did not receive fungicides or insecticides throughout the cropping season. At maturity, the plots of each trial were harvested using a plot combine. From each of the four plots in both trials, three subsamples of 0.9 kg seeds each were collected, resulting in a total of 12 seed samples. All these seed samples were stored at 4 ℃ until use.

### DNA extraction, library preparation, and sequencing

All seed samples were washed with 1% tween 20 for 30s before surface sterilization. The sterilization process involved using 10% bleach for 5 min, followed by 70% ethanol for 30s. The seeds were immediately washed with sterile distilled water thrice and frozen at −20 ℃ until sample processing. DNA from approximately 1g of surface sterilized seeds were isolated using DNeasy Plant Mini Kit (Qiagen, Valencia, CA, USA) with necessary modifications. This included homogenizing five seeds per lysing matrix A (MPbio, OH, USA) using a Bead mill 24 Homogenizer (Thermo Fisher Scientific, CA, USA) at the speed of 4.0 m/s for 30 s, the rest of the protocol were followed based on the manufacturer’s recommendations. DNA integrity was evaluated by electrophoresis on a 1% agarose gel, quality assessed in Quickdrop^TM^ Micro-volume spectrophotometer (Molecular Devices, San Jose, CA, USA), and quantified using a Qubit double-stranded DNA (dsDNA) high-sensitivity assay kit (Thermo Fisher Scientific, MA, USA) in a Qubit 2.0 fluorometer (Invitrogen, CA, USA). Amplicon libraries were prepared by the Genomics and Bioinformatics Service at Texas A&M (College Station, TX, USA) using the primers 515F-GTG CCA GCM GCC GCG GTAA, 806R-GGA CTA CHV GGG TWT CTA AT (Caporaso et al. 2011) targeting the V4 region of 16S rRNA. Similarly, ITS (Internal Transcribed Space)-region based amplicon libraries were prepared using primers ITS1F-CTT GGT CAT TTA GAG GAA GTAA, ITS2-GCT GCG TTC TTC ATC GAT GC (White et al. 1990). Libraries were sequenced on an Illumina Miseq PE250 platform by the Genomics and Bioinformatics Service at Texas A&M University (College Station, Texas, USA). Raw sequences have been deposited in NCBI Sequence Read Archive (SRA) under Bioproject: PRJNA866104 and PRJNA866106 for 16S and ITS amplicon, respectively.

### Sequence processing

All the raw reads were processed using the Mothur v.1.47 (Kozich et al. 2013). The protocols for the Mothur were applied following Miseq sop (https://mothur.org/wiki/miseq_sop/) with few modifications as needed. Total number of reads for were 1,474,695 16S V4 amplicon sequences and 1, 132,133 sequences for ITS-based amplicon after quality filtering. Sequence reads exceeding 400 bp were considered too long for the 16S V4 analysis, while reads below 450 bp were considered too short for the ITS region analysis. In addition, any ambiguous bases and maximum repeat of a nucleotide sequence more than eight were removed using Mothur built-in function “maxambig” and “maxhomop”. One sample from the 16S amplicon was excluded from the analysis due to poor total read quality, resulting in only 11 samples available for processing 16S V4 reads. Mothur built-in function was applied to make contigs, quality control of bad reads, removal of chimeras, and finally assigning operational taxonomic units (OTUs). A 97% cutoff was applied to bacterial taxa, while a 95% cutoff was used for fungal taxa. Ribosomal Database project classifier (Wang et al. 2007) with the Silva database v138 (Quast et al. 2013) was used for taxonomy assignment of 16S rRNA reads. Similarly, Mothur release of UNITE database (Abarenkov et al. 2021) was used for the taxonomic assignments of ITS reads. OTUs assigned to mitochondria, chloroplast, eukaryotes, or unknown were removed from 16S rRNA data sets and OTUs were assigned. Similarly, unknowns were removed from ITS data sets. Before downstream analysis, any single digit OTUs were removed from the dataset to avoid spurious sequences (Reitmeier et al. 2021).

### Diversity estimation and statistical analysis

Relative abundance data from both 16S rRNA and ITS data sets were analyzed in R-studio v2022.2.3.492 (R Studio Team 2022) with R version 4.1.2 (R Core Team 2021) using Phyloseq v1.38 (McMurdie and Holmes 2013), Vegan v2.5.7 (Oksanen et al. 2020), ggplot2 (Wickham 2016), and tidyverse (Wickham et al. 2019). Relative abundance bar graph was generated for both 16S rRNA and ITS data sets, and all taxa with <2% relative abundance were categorized in the “other” category. The alpha-diversity measures were generated for observed, Shannon diversity, Simpson, chao1, and ACE in Phyloseq v1.38 (McMurdie and Holmes 2013). The Chao1 and ACE diversity indices were utilized to assess community richness, whereas the Shannon and Simpson diversity indices were employed to evaluate both richness and evenness within the community (Kim et al. 2017). Mann-Whitney U test of significance was conducted for all alpha diversity measure using “wilcox.test()” built-in function in R. Permutational multivariate analysis of variance (PERMANOVA) was calculated using “adonis()” function in Vegan v2.5.7 (Oksanen et al. 2020) to determine the effects of the two farming systems on the bacterial and fungal communities, with Bray-Curtis dissimilarity matrix on both 16S rRNA and ITS dataset. Non-metric multidimensional scaling (NMDS) plot was constructed based on the Bray-Curtis dissimilarity matrix for visualization. Differentially abundant OTUs were determined in both 16S rRNA and ITS datasets by “lefse” function (Segata et al. 2011) and indicator OTUs were determined using “indicator” function in Mothur v.1.47 (Kozich et al. 2013). Indicator function in Mothur was adapted based on the Dufrene and Legendre (1997) where they defined species to unique to habitats independence of relative abundances. Default parameters were used for both “lefse” and “indicator”.

### Isolation and identification of culturable bacterial endophytes

Approximately 20g of rice seeds were surface sterilized as described above and completely crushed in sterile mortar and pestle with phosphate-buffered saline (PBS) buffer. Crushed rice seeds were then transferred to sterile flask with 200 ml of PBS buffer to create a suspension solution. A serial dilution (100 μl) of the homogenized seed suspension were prepared in PBS buffer and plated on to Tryptic Soy Agar (TSA) plates to determine the culturable population of seed endophytic bacteria. Serial dilutions of 10^-2^, 10^-3^, and 10^-4^ fold were made and 100 μl aliquots of each dilution were spread on separate TSA plates. Colony PCR was conducted to amplify 16S rRNA genes using the universal primers 27F: 5’-AGA GTT TGA TCC TGG CTC AG-3’ and 1492R: 5’-GTT TAC CTT GTT ACG ACT T-3’ (Lane 1991). Amplified PCR products were purified using Zymoclean Gel DNA recovery kits (Zymo Research, CA, USA) using manufacturer’s recommendations. All sequencing were performed by Eton Bioscience (Eton Bioscience Inc., San Diego, CA. USA). Consensus sequences were constructed from the obtained 16S rRNA sequences and identifications were done by Blast in EzTaxon server (https://www.ezbiocloud.net/) (Yoon et al. 2017). The nucleotide sequences of 16S rRNA genes were deposited to the GenBankunder the accession numbers: OP132284-OP132369.

### *In-vitro* bacterial endophyte antagonism assay

Antagonistic activities of seed endophytic bacterial isolates were tested on the three rice seedling blight pathogens: *R. solani* AG4 (Gaire et al. 2020), *R. solani* AG11 (Jones and Carling 2007), and *M. graminum* (Gaire et al. 2021). The isolates were initially dereplicated based on their 16S rRNA gene sequence. Those strains with 97% similarity were grouped together. (Rosselló-Mora and Amann 2001). One representative isolate was selected from each group for the *in-vitro* assay.

The *in-vitro* experiments were conducted using the dual culture method (Huang et al. 2012). All bacterial isolates were grown in LB broth (Thermo Fisher Scientific, Waltham, MA, USA) on an orbital shaker at 150 rpm at 28°C for 48 h. The bacterial suspensions were centrifuged at 4, 032 g for 10 min in 50-ml sterile falcon tubes. The pellets were resuspended with sterile distilled water and the final concentration of all bacterial isolates was adjusted to an OD (optical density) of 0.3 at 600 nm for each isolate. For all fungal pathogen cultures, a 5-mm agar-plug was taken from the edge of the colony of actively growing fungal cultures on Potato Dextrose Agar (PDA) plates and placed at the center of another PDA plate. Each of the four locations, equally spaced at a distance of 2 cm from the central fungal culture disk, received a drop (10 μl) of bacterial suspension. Sterile distilled water was used as the negative control. The test plates were incubated at 27°C. Each treatment was replicated three times, and the experiments were repeated twice. The mycelia growth (cm) of the three-fungal pathogens, *R. solani* AG4, *R. solani* AG11, and *M. graminum*, was measured at different time points: 2 days, 3days, and 5 days post incubation, respectively, considering their distinct growth rates.

### Phylogenetic analyses

Phylogenetic analyses were performed using consensus 16S rRNA sequences of endophytic bacterial isolates that exhibited antagonistic activity against rice seedling blight pathogens. For each of the bacterial isolates, top hits in the databases were compared to generate a phylogenetic tree. All sequences were aligned with Silva database v138 (Quast et al. 2013) and alignment were edited and adjusted using Mothur v1.47 (Kozich et al. 2013). On all the aligned sequences Neighbor-Joining (NJ) (Saitou and Nei 1987) phylogenetic analyses were performed in R-studio with R version 4.1.2 using APE package version 5.4-1 (Paradis et al. 2004). All trees were visualized and manually edited using FigTree v1.4.4 (http://tree.bio.ed.ac.uk/software/figtree/) (Rambaut 2018).

## RESULTS

### Microbial communities associated with rice seed endosphere

A combined total of 1,474,695 sequences were obtained for 16S rRNA V4 region sequencing, while 1,132,133 sequences were acquired for ITS amplicon sequencing. One sample from 16S rRNA sequences was excluded from analysis due to extremely low-quality sequences. Analysis of 16S rRNA amplicons identified a total of 78 OTUs after removal of single digit OTUs. The predominant bacterial orders within the core community across all the seed samples were *Enterobacterales*: 46.38 to 57.39%, followed by *Rhizobiales*:11.48 to 16.08%, *Micrococcales:* 8.35 to 12.19%, *Xanthomonadales*: 5.85 to 9.25%, *Pseudomonadales*: 5.45 to 7.54%, *Sphingomonadales*: 3.05 to 4.63%, *Paenibacialles*: 1.71 to 2.60%, *Burkholderiales*: 0.44 to 4.17%, *Kineosporiales*: 0.69 to 2.71%, and other orders with less than 2% on all the samples were placed in “others” category (Fig. 1; Supplementary Table 1). These nine orders were classified into 11 different genera, where Unclassified *Enterobacterales* dominated the genera classification among both treatments.

**Fig. 1.**
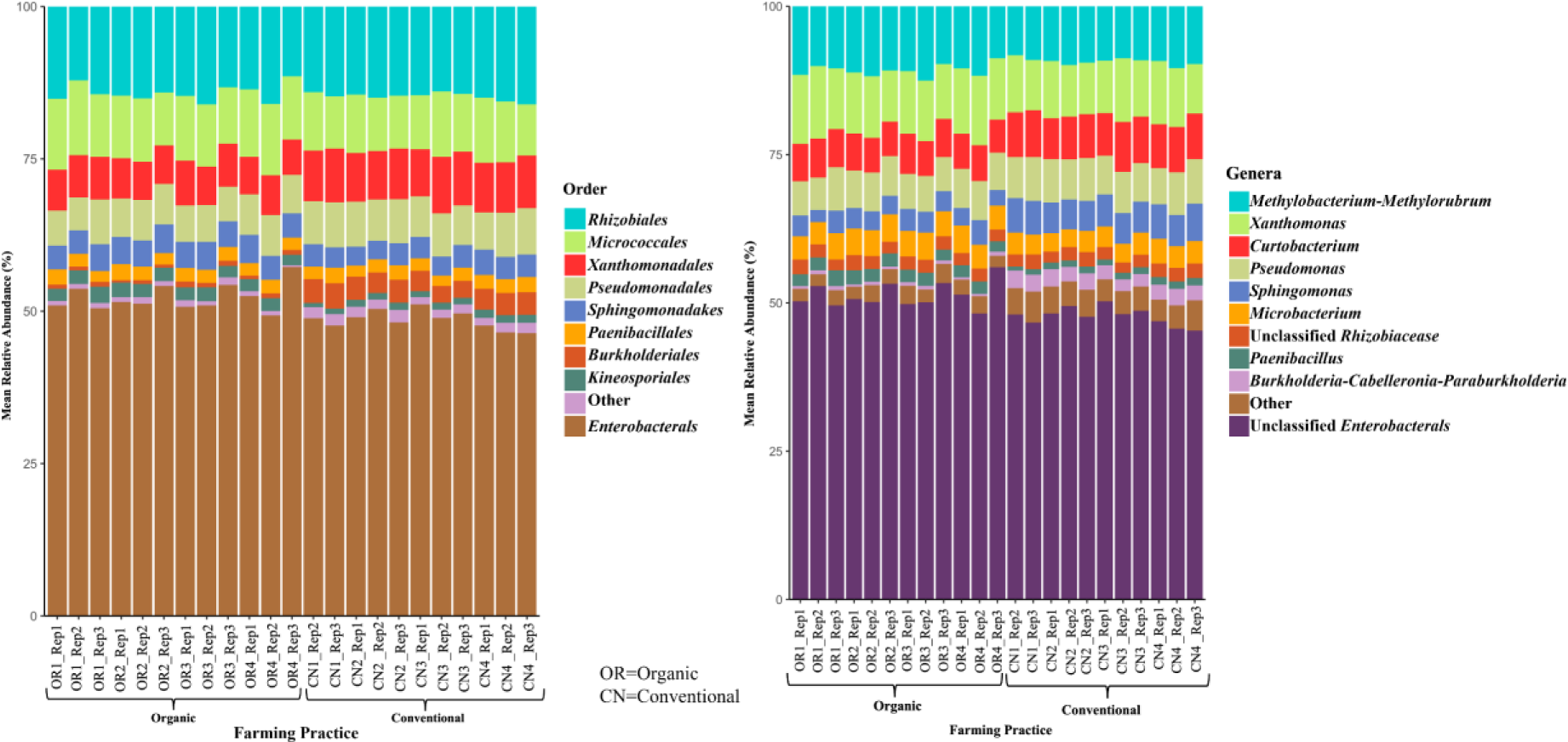
Relative abundance of seed endophytic bacterial populations in rice grown in organic and conventional farming systems based on 16S rRNA V4 region amplicon sequencing. Bacterial taxa with less than 2% relative abundance are placed in the ‘other’ category. OR represents the seed samples collected from organic rice whereas CN represents the seed samples from conventional rice.

Similarly, ITS amplicons sequencing analysis identified a total of 54 OTUs after removal of single digit OTUs. The *Pleosporales* were the dominant fungal order, accounting for >90% relative abundance in all samples (Fig. 2; Supplementary Table 2). When classifying the OTUs at the genera level, we observed differences in the abundance of *Phoma,* unclassified *Pleosporales,* and *Cochliobolus* between the farming practice treatments. In conventional rice seeds, the fungal genus *Phoma* was the dominant fungal genera, ranging from 49.08 to 55.98%, while in organic rice seeds, it ranged from 9.18 to 14.34%. In organic rice seeds, the unclassified *Pleosporales* were the dominant genera, ranging from 38.33 to 41.01%, while in conventional rice seeds, it ranged from 19.7 to 24.09%. In addition, organic rice seeds exhibited a higher abundance of fungal genus *Cochliobolus* compared to conventional rice seeds, ranging from 4.36 to 13.04% and 0.57% to 3.60, respectively.

**Fig. 2.**
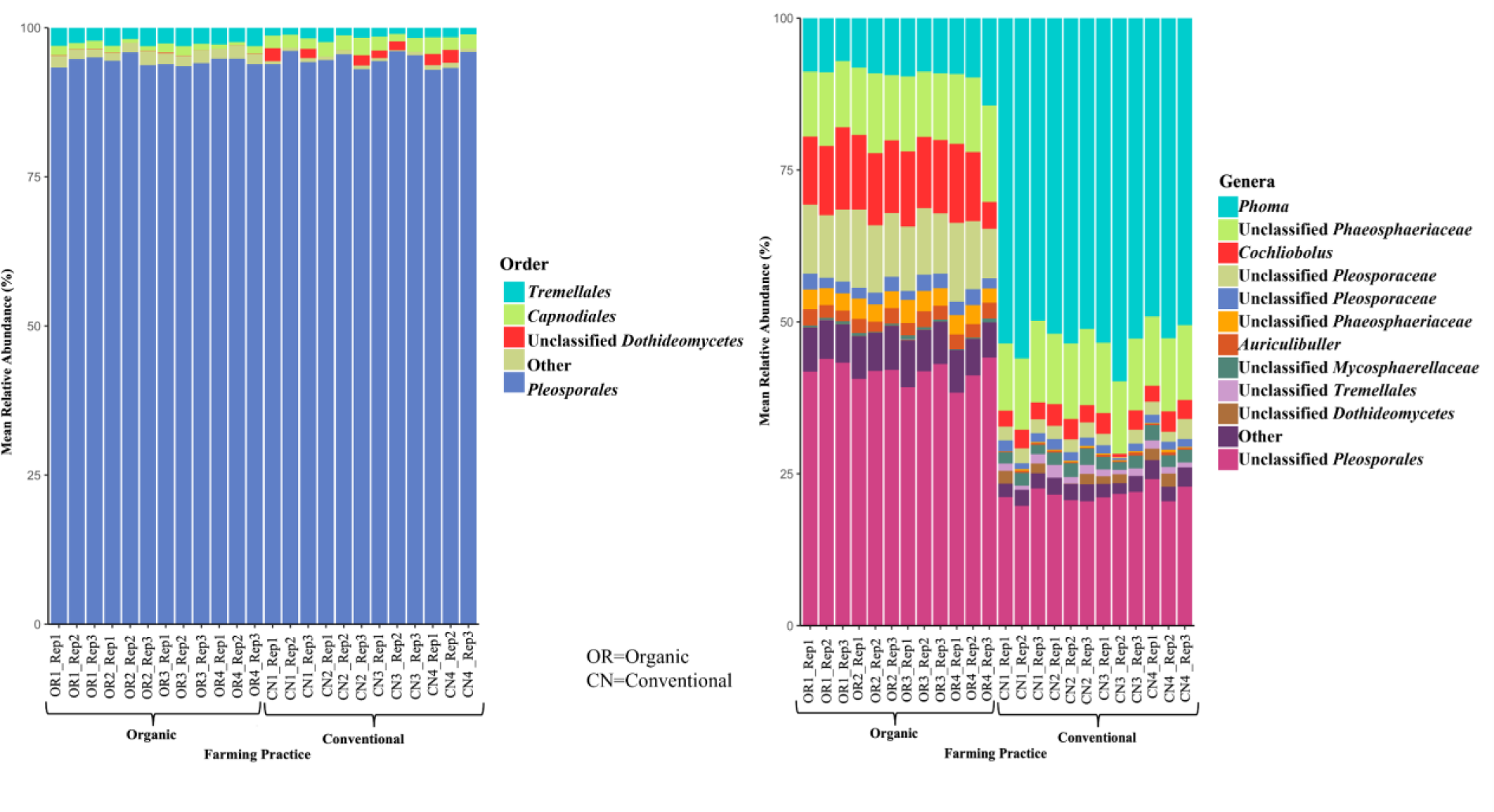
Relative abundance of seed endophytic fungal populations in rice grown in organic and conventional farming systems based on ITS-amplicon sequencing. Fungal taxa with less than 2% relative abundance are placed in the ‘other’ category. OR represents the seed samples collected from organic rice whereas CN represents the seed samples from conventional rice.

### Variation in microbial diversity between conventional and organic rice seeds

The nonmetric multidimensional scaling (NMDS) ordination of variations in bacterial and fungal communities showed the strong clustering of endophytic microbial assemblage among the seeds obtained from conventional and organic farming systems (Fig. 3). The observed difference between the two farming systems based on the Bray-Curtis distance was significant.

**Fig. 3.**
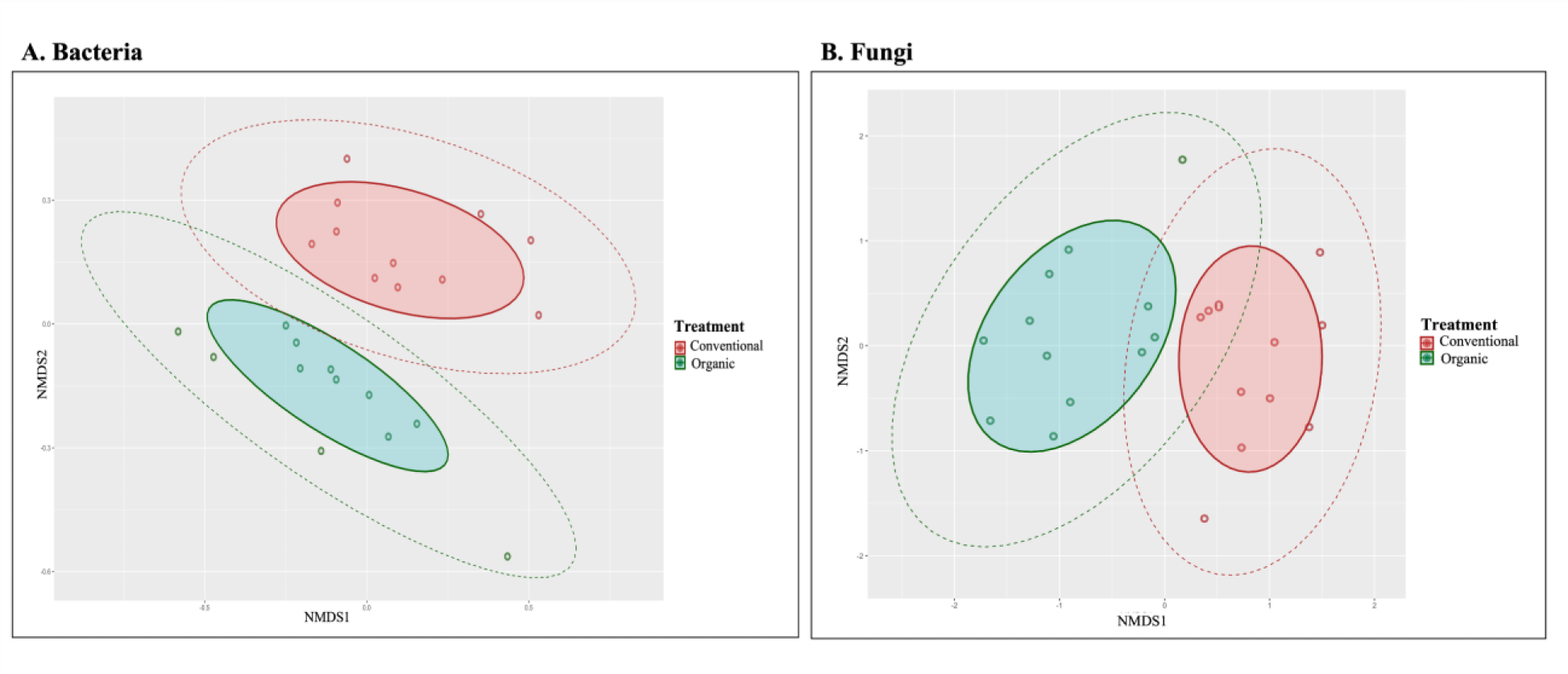
Nonmetric multidimensional scaling (NMDS) ordination of variation in bacterial (A) and fungal (B) community structure (Bray-Curtis distance) in the endophytic microbial populations of rice seeds grown in organic and conventional farming systems. PERMANOVA based on Bray-Curtis distance, bacteria=0.001(*R^2^*=0.719), fungi=0.001 (*R^2^*=0.964) Shaded area represents 75% of the core composition whereas dashed line represents 95% of the composition.

The result of PERMANOVA on the Bray-Curtis distance showed that the difference in farming system explained 71.9%(*p*<0.001) of the variation in the bacterial community whereas in fungal community the same analysis explained 96.4% (*p*<0.001) of the variation between two farming systems.

Similarly, an analysis of alpha diversity measured was performed on observed, Shannon, and Simpson indices. We used Mann-Whitney U test to analyze the alpha diversity measures. Our analysis on the bacterial community (Fig. 4A) showed among community richness only ACE estimator ACE (W=99, df = 1, *p* < 0.05) was significantly different whereas, there was no significant difference in observed/sobs (W=95, df = 1, *p* = 0.07831), and other community richness indices Chao1 (W=94, df = 1, *p* = 0.09039) between bacterial communities in the conventional and organic rice seeds. In contrast, both indices Shannon (W=127, df = 1, *p* < 0.0001) and Simpson (W=8, df = 1, *p* < 0.0001) were significantly higher in conventional rice seeds in comparison to organic rice seeds. Analysis of diversity indices on fungal community (Fig. 4B) showed significant higher diversity on community richness indices, observed/sobs (W= 107, df = 1, *p* < 0.05) and community richness indices in Chao1 (W=112, df = 1, *p* < 0.02205), and both Shannon (W=0, df = 1, *p*< 0.0001) and Simpson W=144, df = 1, *p* < 0.0001) indices on organic rice seeds in comparison to conventional rice seeds. Only one community richness index, ACE (W=95, df = 1, *p* = 0.1978) showed no significant differences between fungal communities in organic rice seeds compared to conventional rice seeds.

**Fig. 4.**
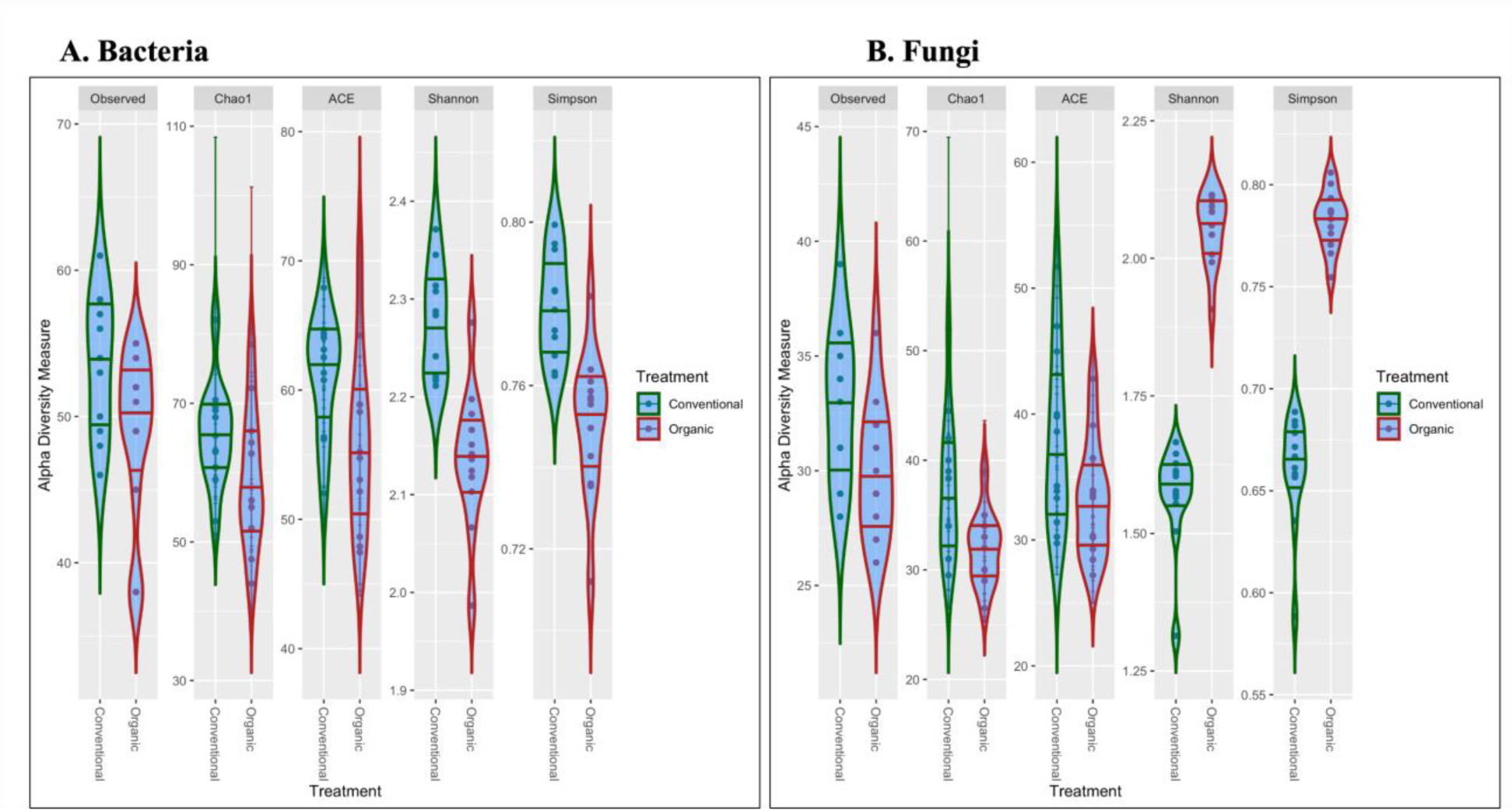
Diversity indices (observed, chao, ACE, Shannon, and Simpson) of bacterial (A) and fungal (B) communities in endophytic microbial populations in rice seeds grown in organic and conventional farming systems. Each point in the violin-blot indicates the individual sample. A, Observed (W=95, df = 1, *p* = 0.07831), Chao1 (W=94, df = 1, *p* = 0.09039), ACE (W=99, df = 1, *p* < 0.05), Shannon (W=127, df = 1, *p*< 0.0001), and Simpson (W=8, df = 1, *p* < 0.0001). B, Observed (W= 107, df = 1, *p* < 0.05), Chao1 (W=112, df = 1, *p* < 0.02205), ACE (W=95, df = 1, *p* = 0.1978), Shannon (W=0, df = 1, *p*< 0.0001), and Simpson W=144, df = 1, *p* < 0.0001).

### Drivers of variation in community composition and diversity

The Linear Discriminatory Analysis (LDA) effect size (Lefse) was conducted on both bacterial (Fig. 5A) and fungal (Fig. 5B) communities. Our analysis revealed the differentially abundant OTUs between the conventional and organic rice seeds. Of the 36 OTUs that were differentially abundant taxa among the bacterial OTUs, 19 were differentially abundant in conventional rice seeds while the rest of 17 OTUs in organic rice seeds (Fig 5A). In addition, we also analyzed the indicator OTUs, where we define indicators as specific OTUs present across all the samples of one single treatment. Our analysis revealed 12 indicator OTUs in the rice seeds from conventional farming system: Otu009 (genera: *Methylobacterium*, order: *Rhizobiales*), Otu017 (genera: *Luteibacter*, order: *Xanthomonadales*), Otu022 (genera: Unclassified *Comamonadaceae*, order: *Burkholderiales*), Otu024 (genera: *Mucilaginibacter*, order: *Sphingobacteriales*), Otu029 (genera: *Sphingomonas*, order: *Sphingomonadales*), Otu030 (genera: *Williamsia*, order: *Corynebacteriales*), Otu038 (genera: Unclassified *Microbacteriaceae*, order: *Micrococcales*), Otu048 (genera: *Stenotrophomonas*, order: *Xanthomonadales*), Otu052 (Unclassified *Saccharimonadales*), Otu058 (genera: *Bosea*, order: *Rhizobiales*), Otu062 (genera: *Novosphingobium*, order: *Sphingomonadales*), and Otu067 (genera: *Aeromicrobium*, order: *Propionibacteriales*). Nine out of 12 OTUs overlapped with differentially abundant OTUs in conventional rice (Fig. 5A). We did not find any indicator OTUs in rice seeds from the organic farming system.

**Fig. 5.**
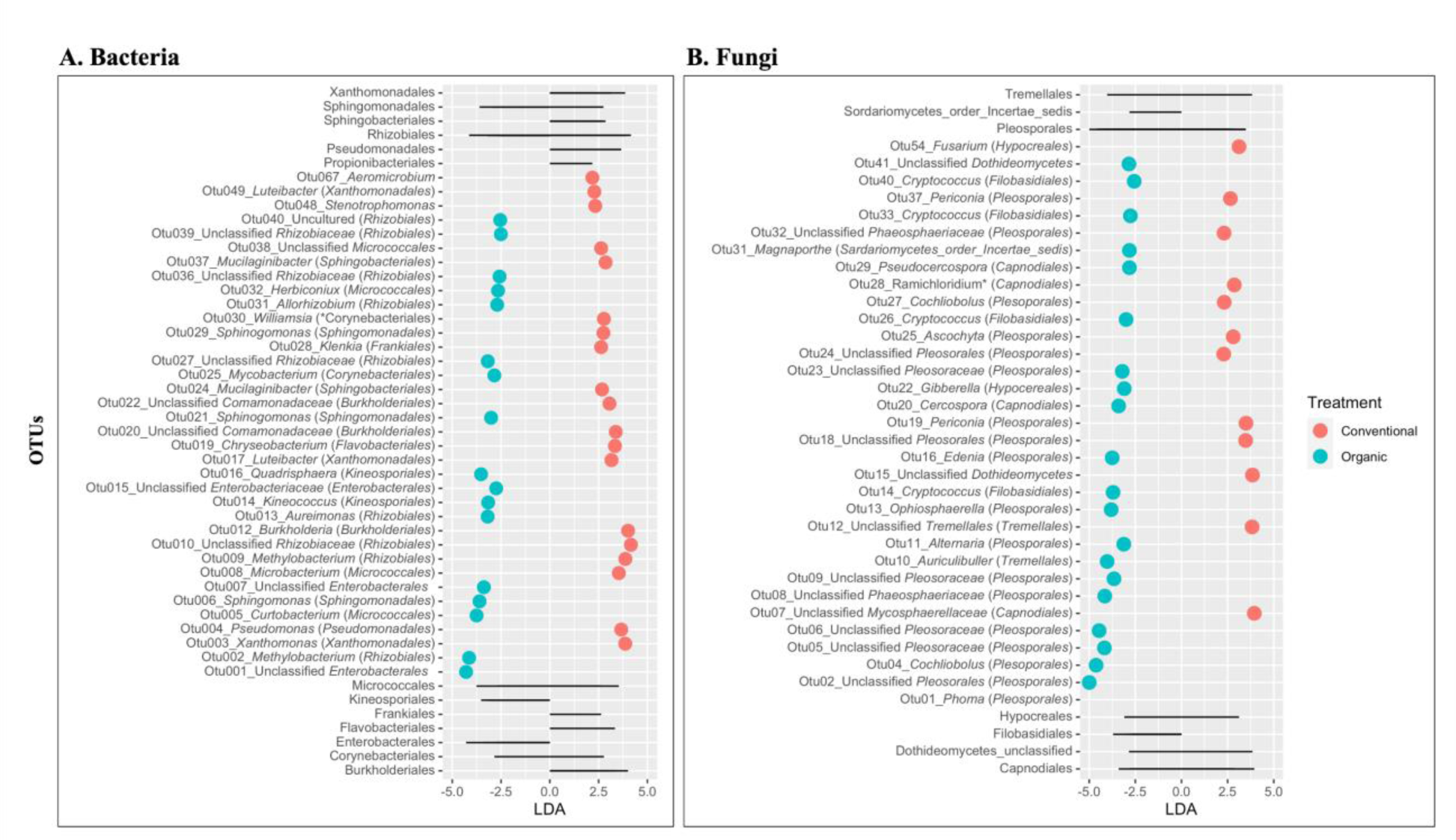
Least Discriminatory Analysis (LDA) effect size (Lefse) on bacterial (A) and fungal (B) communities of the endophytic microbial populations in rice seeds grown in conventional and organic farming systems. Significance was determined using default parameters (Kruskal Wallis test *p* <0.05 and LDA score >2). Each point represents the magnitude of effect size by specific OTUs in individual treatments. Gray lines on top and bottom indicate the orders of OTUs that are differentially abundant in both bacterial and fungal communities.

Similarly, Lefse analysis on fungal community (Fig 5B) revealed 34 differentially abundant OTUs. Of these, 13 OTUs were differentially abundant in rice seeds from the conventional farming system and 21 OTUs were in rice seeds from the organic farming system. Our indicator analysis revealed seven indicator OTUs in rice seeds from the conventional farming system: Otu01 (genera: *Phoma*, order: *Pleosporales*), Otu07 (genera: Unclassified *Mycosphaerellaceae*, order: *Capnodiales*), Otu015 (Unclassified *Dothideomycetes*), Otu018 (Unclassified *Pleosporiales*), Otu028 (genera: *Ramichloridium*, order: *Capnodiales*), Otu032 (Unclassified *Phaeosphaeriacea,* order: *Pleosporales*), Otu037 (genera: *Periconia*, order: Pleosporales), and Otu054 (genera: *Fusarium,* order: *Hypocereales*). All seven-indicator fungal OTUs overlapped with 13 OTUs differentially abundant in the conventional farming systems (Fig. 5B).

### Culture-dependent bacterial diversity

Bacterial endophytes were isolated from rice seeds grown in the conventional and organic farming systems. Titer counts of seed endophytic bacteria were 6 x 10^8^ cfu/ml and 7 x 10^8^ cfu/ml for conventional and organic rice seeds, respectively. A total of 39 and 46 bacterial endophytes were isolated and purified from conventional and organic rice seeds, respectively (Table 1). All 85 bacterial isolates were subjected to near full-length 16S rRNA sequencing for identification. Bacteria that belonged to genera *Pantoea* and *Pseudomonas* dominated the seed endosphere in both farming systems. *Pantoea* isolates accounted for 41% of the total bacterial seed endophytic community in conventional rice seeds and 33% in organic rice seeds. Similarly, *Pseudomonas* spp. accounted for 20% and 21% of the total seed endophytic bacteria in conventional and organic rice seeds, respectively. Among the rare isolates, *Curtobacterium* spp. accounted for 5% of seed endophytic bacterial isolates in conventional rice seeds whereas they covered 13% of seed endophytic bacterial isolates in organic rice seeds. Furthermore, two *Bacillus* spp. were identified in conventional rice seeds and one each of *Paenibacillus* spp. and *Chryseobacterium* spp. identified in organic rice seeds (Table 1).

**Table 1.**
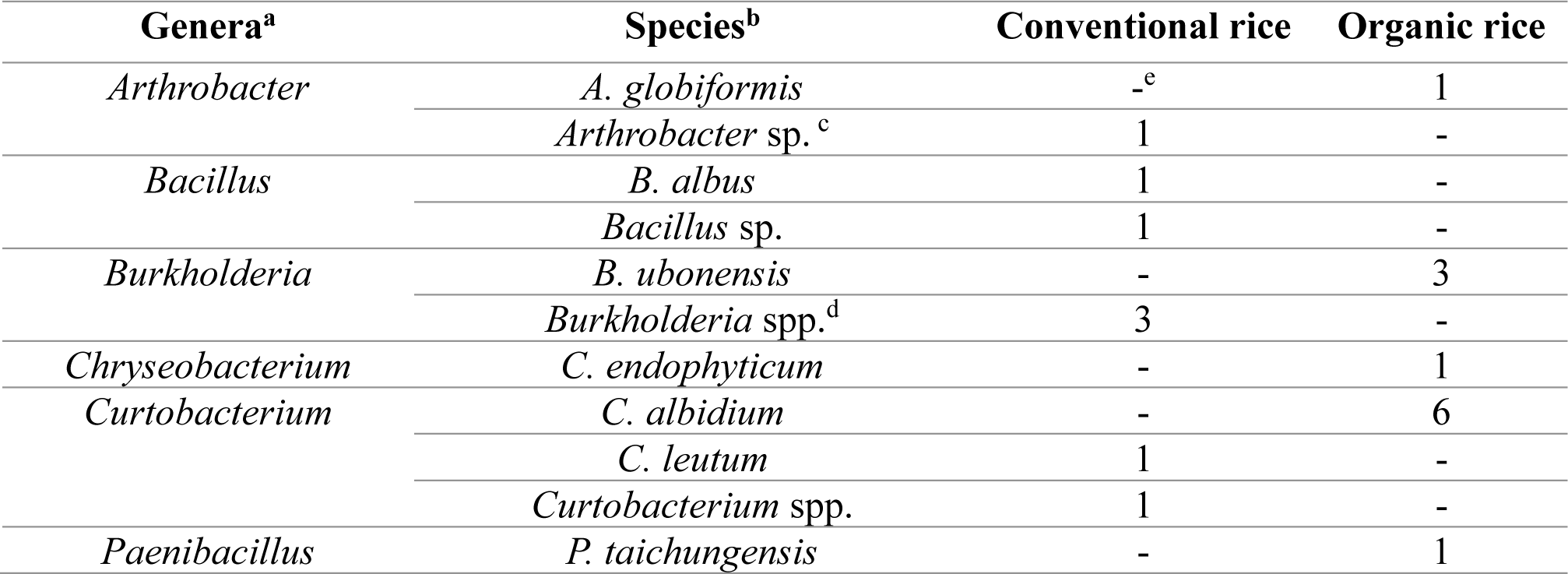
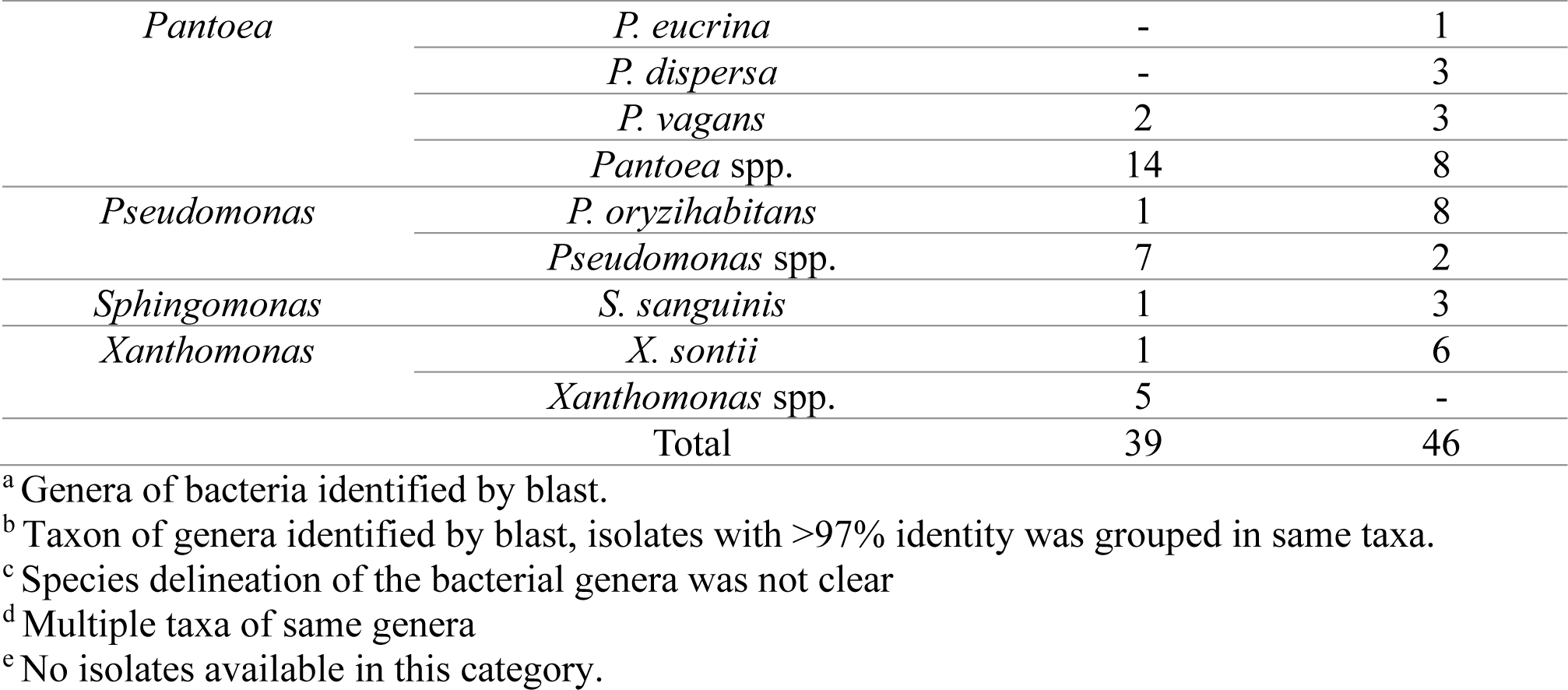
Identification of culturable seed endophytic bacteria isolated from rice seeds grown in conventional and organic farming systems.

### Antagonistic activity of bacterial endophytes against seedling blight pathogens

From the endophytic bacteria isolated from both conventional and organic rice seeds 31 representative isolates were randomly selected based on their 16S rRNA sequences differences. Isolates with >97% sequence similarities were grouped together as putative same “species” and a representative isolate was selected from each group for the experiment. Out of the 31 isolates selected, 11 were derived from organic rice seeds while the remaining 20 were isolated from conventional rice seeds. *In-vitro* assays were conducted with the 31 different bacterial isolates to evaluate their antagonistic activity against three seedling blight in rice pathogens: *M. graminum, R. solani* AG4, and *R. solani* AG11. Three isolates: *Bacillus* sp. ST24, *Burkholderia* sp. OR5, and *Pantoea* sp. ST25. exhibited antagonistic properties against the three seedling blight pathogens. (Table 2, Supplementary Fig. 1). These three bacterial isolates demonstrated the ability to inhibit more than 50% of all three seedling blight pathogens. On the contrary, two *Burkholderia* isolates ST35 and ST43, showed radial enhancement in *R. solani* AG4. (Table 2). Additionally, the phylogeny analysis was performed on three bacterial species, *Bacillus*, *Burkholderia*, and *Pantoea*, using 16S rRNA of each sequences available for each species in databases NCBI nucleotide (https://www.ncbi.nlm.nih.gov/nuccore) and RDP (http://rdp.cme.msu.edu/classifier/classifier.jsp). The phylogenetic analyses conducted for all three bacteria species resulted in their separate classification within distinct clades (Supplementary Fig. 2).

**Table 2.**
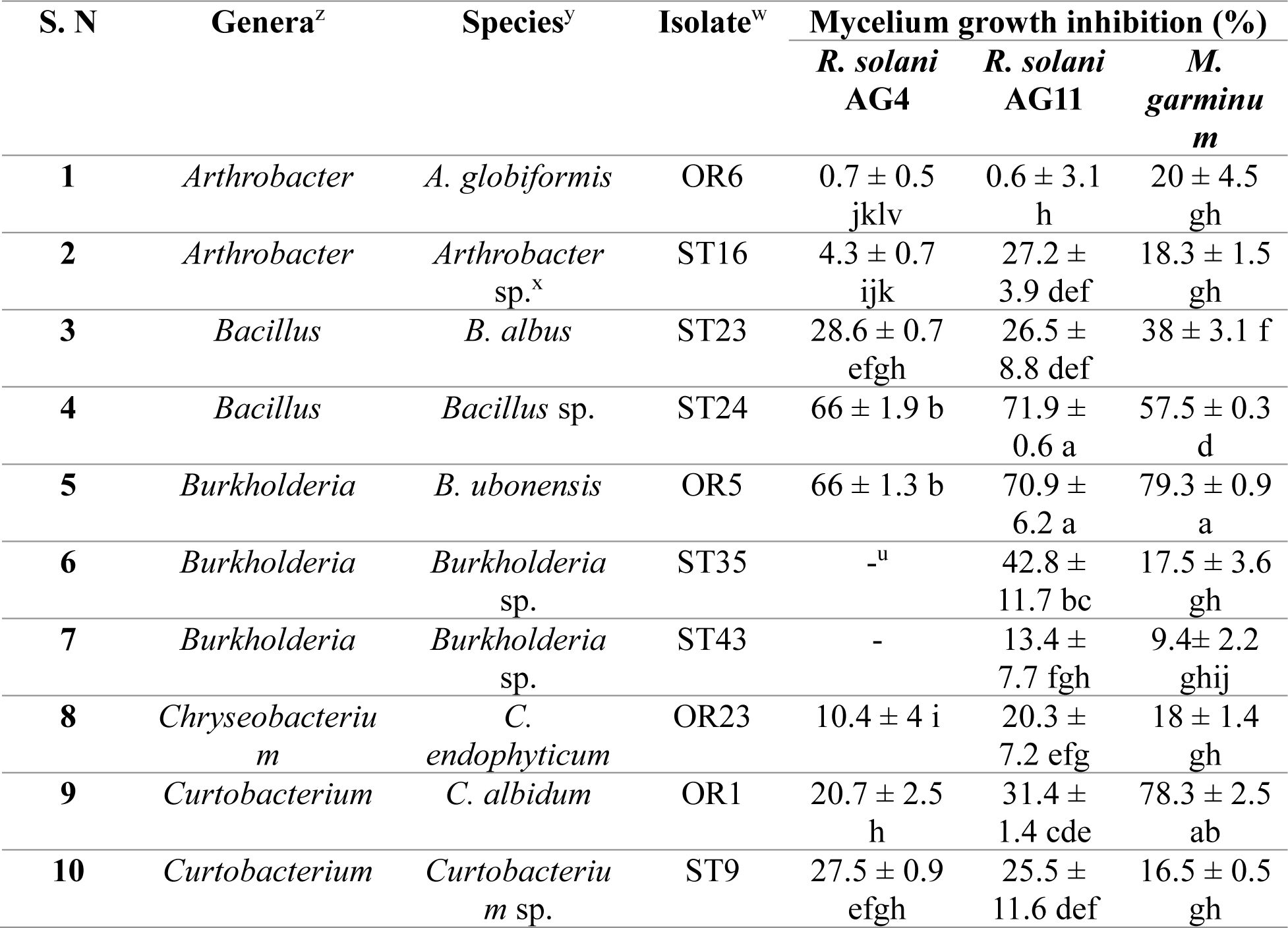
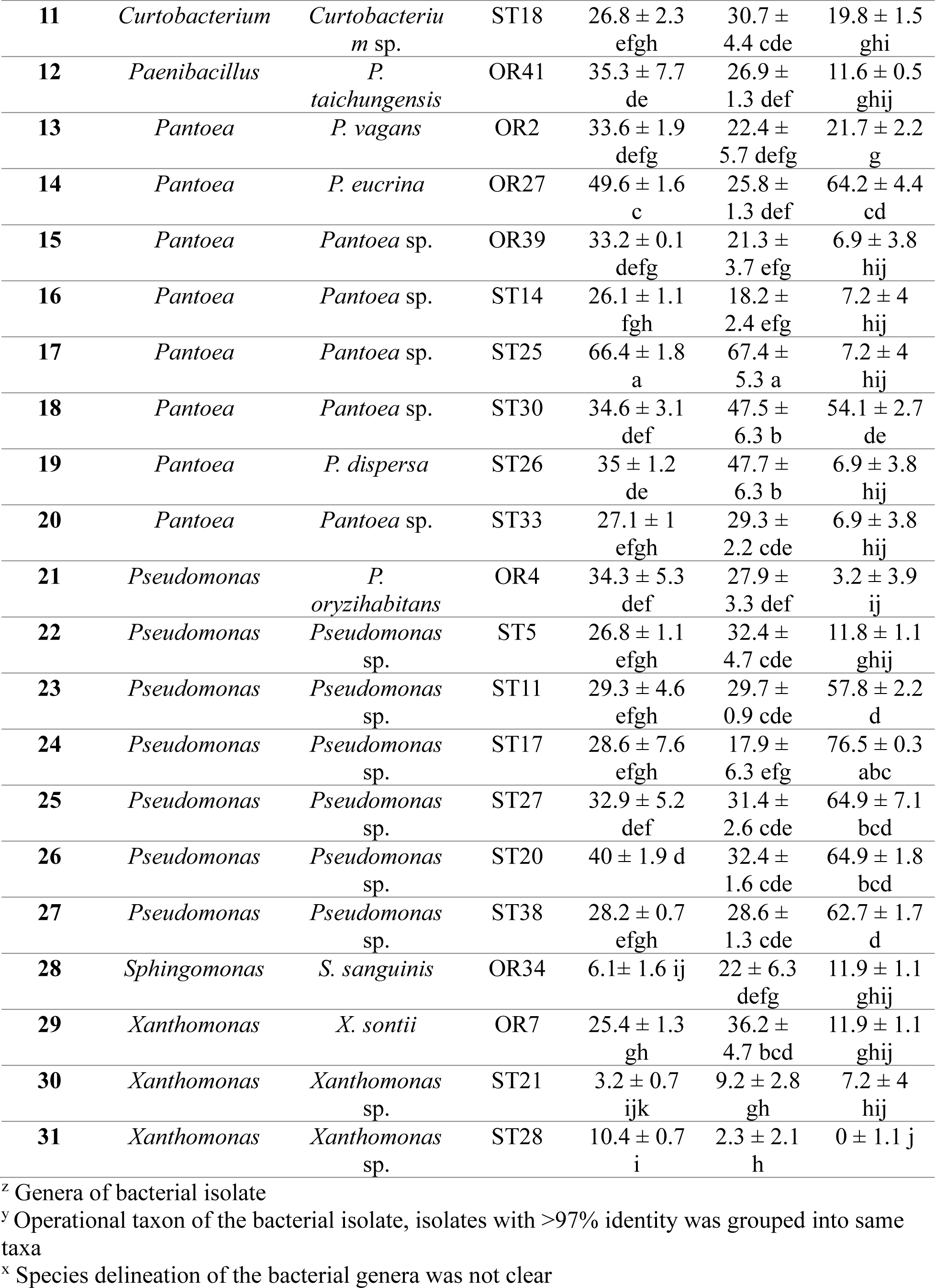
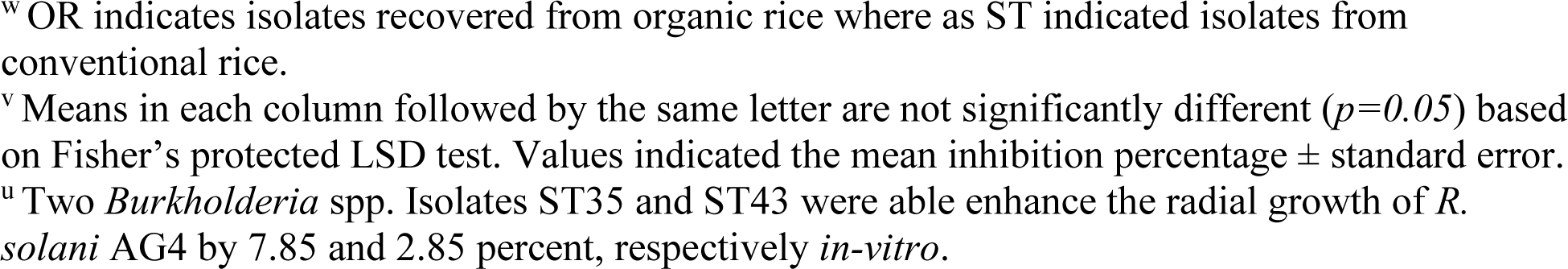
Identification of seed endophytic bacteria and their *in-vitro* antagonistic activities against the three rice seedling disease pathogens, *Marasmius graminum, Rhizoctonia solani* AG4 and *R. solani* AG11.

## DISCUSSION

The plant endophytes residing within seeds (spermosphere) hold a significant role in the plant’s life cycle. They contribute to the maintenance of microbial populations across successive plant generations, making them integral to the plant’s life cycle. In addition to providing beneficial endosymbionts to the offspring, they also actively contribute to seed conservation, facilitate seed germination in soil, and enhance plant growth and defense mechanisms (Cope-Selby et al. 2017; Nelson 2018; Shade et al. 2017; Shahzad et al. 2018). Horizontal transmission, alongside vertical transmission, contributes to the overall composition of the seed microbiota (Shade et al. 2017; Nelson 2018). Previous studies have shown varying effects of farming practices on soil and root microbial communities in different cropping systems (Armalytė et al. 2019; Gonthier et al. 2014; Harkes et al. 2019; Hartmann et al. 2015; Liu et al. 2007; Lori et al. 2017; Lupatini et al. 2017; Sinong et al. 2021). In this study, we employed both culture-independent and culture-dependent approaches to evaluate the endophytic microbial communities within rice seeds produced under conventional and organic farming systems.

The results of the current study demonstrate, for the first time, the differences in seed endophytic microbial communities between rice grown in conventional and organic farming systems. Eyre et al. (2019) have identified and characterized the core rice seed microbiome. However, their results are limited to conventional rice farming systems only since their research was based on rice seeds harvested from rice crops grown under conventional management. In the current study, we compared the differences in the core microbiome of rice seeds between conventional and organic production systems. Although the observed endophytic bacterial communities were not significantly different between the two farming systems evaluated, alpha diversity measures revealed the higher diversity of bacterial communities in the conventional farming system. Bacterial communities in the seed have been observed to be highly variable and they are not only dependent on agricultural practices, but host plant species as well (Tkalec et al. 2022). Therefore, more investigations are needed to further understand if the observed diversity between organic and conventional rice is consistent across the genotypes or restricted to one. From the current study, *Proteobacteria, Actinobacteriota,* and *Bacteriota* were the most abundant phyla, which is consistent with the rice seed core microbiome reported before (Eyre et al. 2019). Additionally, in our study, the highest observed genera were the unclassified *Enterobacterales.* However, these findings do not reveal a distinct pattern between the two farming systems. Some genera such as *Curtobacterium* and *Methylobacterium* were observed consistently in rice seeds grown under organic management; similarly, *Burkholderia* was observed at a higher abundance in rice seeds from the conventional management system. Of these taxa it is noteworthy that methylotrophic bacteria (such as *Methylobacterium)* are well known growth promoters for various crop plants (Kumar et al. 2019). All these observed taxa have been associated with the rice seed microbiome reported previously (Eyre et al. 2019). Furthermore, these bacterial taxa have also been detected in soil microbiomes of both conventional and organic farming systems for various crops such as wheat, barley, maize, and other non-rice crops (Lupatini et al. 2017). Nonetheless, the transfer of soil microbiota to seed microbiota does not always occur directly as various factors involve during the establishment of seed microbial populations (Nelson 2018; Shade et al. 2017). Among the endophytic fungal communities, the ITS-region based amplicon sequence data reveals higher diversity of fungal communities in rice seeds grown under the organic farming system. The higher fungal diversity in organic systems over the conventional system can be explained by the impacts of the use of synthesized chemicals, especially fungicides, in the conventional farming system. No synthesized chemicals were used in the organic farming systems, whereas different synthesized chemicals such as fungicides, insecticides, and herbicides were applied in the conventional farming system. It has been well known that pesticides, especially fungicides, can selectively inhibit or eliminate certain groups of fungal and other microbial populations (Liu et al. 2007; Lupatini et al. 2017). In the current study, *Ascomycota* and *Basidiomycota* were only two phyla observed in rice seeds, which is consistent with the results of the rice core microbiome study reported previously (Eyre et al. 2019). In contrast to the endophytic bacterial communities, we observed a clear taxonomic pattern within the fungal communities. Approximately 50% of the total genera in rice seeds cultivated under the conventional farming practices consisted of *Phoma* spp. These *Phoma* spp. were widely distributed and known as phytopathogens, sometimes associated with food contaminants with clinical significance, especially in immunocompromised individuals (Bennett et al. 2018). Among the fungal communities in rice seeds grown in the organic farming system, unclassified *Pleosporales* dominated the fungal taxa. We also observed a higher abundance of *Cochliobolus*spp. in rice seeds grown in the organic farming system. Certain strains belonging to *Cochliobolus* genera have been identified as established pathogens of rice (Marchetti and Peterson 1984). The current study reveals that the reduced occurance of unclassified *Pleosporales* and *Cochliobolus* in conventional seeds, compared to organic seeds, may be attributed to the application of fungicides such as azooxystrobin and propiconazole on conventional rice cultivation. These fungicides likely inhibited the presence of these fungi. However, ITS does not provide full resolution for definitive species identification. Furthermore, *Auriculibuller* spp. was also observed in rice seeds grown in the organic farming systems, which has been previously reported as one of the compositions of aerial microbiota present in rice fields (Franco Ortega et al. 2020). The usage of additional locus or metagenomic surveys in future studies can provide more in-depth understanding of the diversity and functional implication of seed endophytic communities in rice grown under conventional and organic farming systems. When studying the plant endosphere, the host genome overwhelms the genetic composition within the total metagenomes. To gain deeper insights into seed communities, new techniques such as the “microbial bait” approach can be employed to capture and analyze the microbiota.

In our culture-dependent microbial survey, no difference was observed in the endophytic bacteria between rice plants grown under conventional and organic management systems. All the isolated seed endophytic bacteria were classified into 10 different genera. Notably, the bacterial groups *Panotea, Pseudomonas,* and *Xanthomonas* were isolated at a higher percentage, consistent with previously observed endophytic bacterial groups found in rice seeds (Hardoim et al. 2012; Mano and Morisaki 2008; Mano et al. 2006; Walitang et al. 2017). In this study, we identified other endophytic bacterial genera including *Arthrobacter, Bacillus, Curtobacterium, Chryseobacterium, Paenibacillus,* and *Sphingomonas.* These genera have also been previously isolated as rice endophytic bacteria (Mano and Morisaki 2008; Mano et al. 2006). Many of these isolated bacterial genera are known to establish positive symbiotic association with plants, exhibiting various plant growth promotion activities and providing protection against phytopathogens. We evaluated the ability of these seed endophytic bacterial isolates to protect rice seed from the rice seedling blight pathogens. Our study demonstrated that two strains, one belonging to *Bacillus* spp. and other to *Pantoea* spp., exhibited *in-vitro* suppression of the three rice seedling blight pathogens, *R. solani* AG4, *R. solani* AG11, and *M. graminum*. *Bacillus* spp. and the other *Pantoea* spp. are well known for their pathogen-suppressing abilities (Asaka and Shoda 1996; Huang et al. 2012; Jamali et al. 2020; Ritpitakphong et al. 2016; Walterson and Stavrinides 2015; Wu et al. 2019). *Burkholderia* spp. isolates were observed to enhance the radial growth of *R. solani* AG4. Bacterial-fungal symbiosis plays an important role in mutualistic growth and improved fitness of both bacterial and fungi (Abeysinghe et al. 2020; Lackner et al. 2009), has been reported previously in both *Rhizoctonia* (Obasa et al. 2017) and *Burkholderia* (Lackner et al. 2009). However, further research is required to gain a better understanding of the ability of these bacterial isolates to protect rice seeds against seedling blight pathogens, as well as potential role of these fungal symbionts in disease development and pathogenesis. Furthermore, these seed endophytic bacteria may possess several plant growth-promoting activities. However, characterization of other plant growth promoting activities exhibited by all isolated bacterial genera in this study is beyond the scope of our research.

The present study represents a proof of concept with limitations in terms of temporal sampling (single time point) and spatial extent (local) for evaluating the differences in microbial diversity within seeds between conventional and organic farming practices. Further studies are needed to establish a more comprehensive framework that can provide a deeper understanding of seed endophytes and their associations with farming practice. We also underline the necessity of long-term studies, encompassing diverse ecosystems relevant to rice agriculture and incorporating different management systems and geographic scales, to better understand the complexities of the farming systems and seed microbiota.

Additionally, we acknowledge the limitations of the culture-dependent approach employed in this study. Regardless, our study has provided valuable insights into the potential of these seed endophytic microbial populations. This highlights the significance of culture-based studies in assessing the functional capabilities of the observed microbiome differences. Additionally, the lack of taxonomic resolution in our culture-dependent approach could be attributed to the use of limited culture conditions. Emerging high throughput culturomics methods, despite being costly, hold promise for future investigations, offering a more comprehensive understanding of the functional potential of seed endophytes, especially in relation to farming practices. These microbial populations have potential applications in modern agriculture, where scientists and growers constantly strive to adapt to changing climatic conditions, diminishing effectiveness of chemical-based protections, and evolving dynamics of phytopathogens.

In summary, we, for the first time, identified and compared the bacterial and fungal microbial communities within rice seeds produced under conventional and organic farming systems. Our findings reveal distinct differences in the seed microbial populations between both farming practices The bacterial endophyte populations in conventional rice seeds exhibited higher diversity compared to those in organic rice seeds, whereas the fungal endophyte populations showed greater diversity in organic rice seeds compared to conventional rice seeds. Additionally, we identified bacterial endophytes with potential as biological control agents against three seedling-blight pathogens. Overall, our results contribute new insights into the structure and composition of seed endophytic microbial communities, as well as the influence of farming practice (conventional vs. organic) on these communities. This knowledge can help develop novel microbiome-based approaches, such as the creation of biotic stress tolerant crops, to enhance plant health and improve crop productivity.

## Author Contributions

S. K., X-G., Z., and S. A. B conceptualized the project. S. K and M. I. worked with culturable isolates collection and antagonistic ability experiments. S. K. performed bioinformatics analyses and all other data analysis. S. K., X-G., Z., and S. A. B. drafted the manuscript, and all authors contributed to the final version. X.-G. Z. provided funding to the research.

## ACKNOWLEDGEMENTS

Portions of this research were conducted with the advanced computing resources provided by Texas A&M High Performance Research Computing. This work was supported, in part, by the Texas Rice Research Foundation (Grant nos. M2101762 and M2202432) and USDA NIFA OREI (2015-51300-24286).

