## Supplementary figures and images for "Characterization of Differences of Seed Endophytic Microbiome in Conventional and Organic Rice by Amplicon-based Sequencing and Culturing Methods"

### Supplemental Figure 1

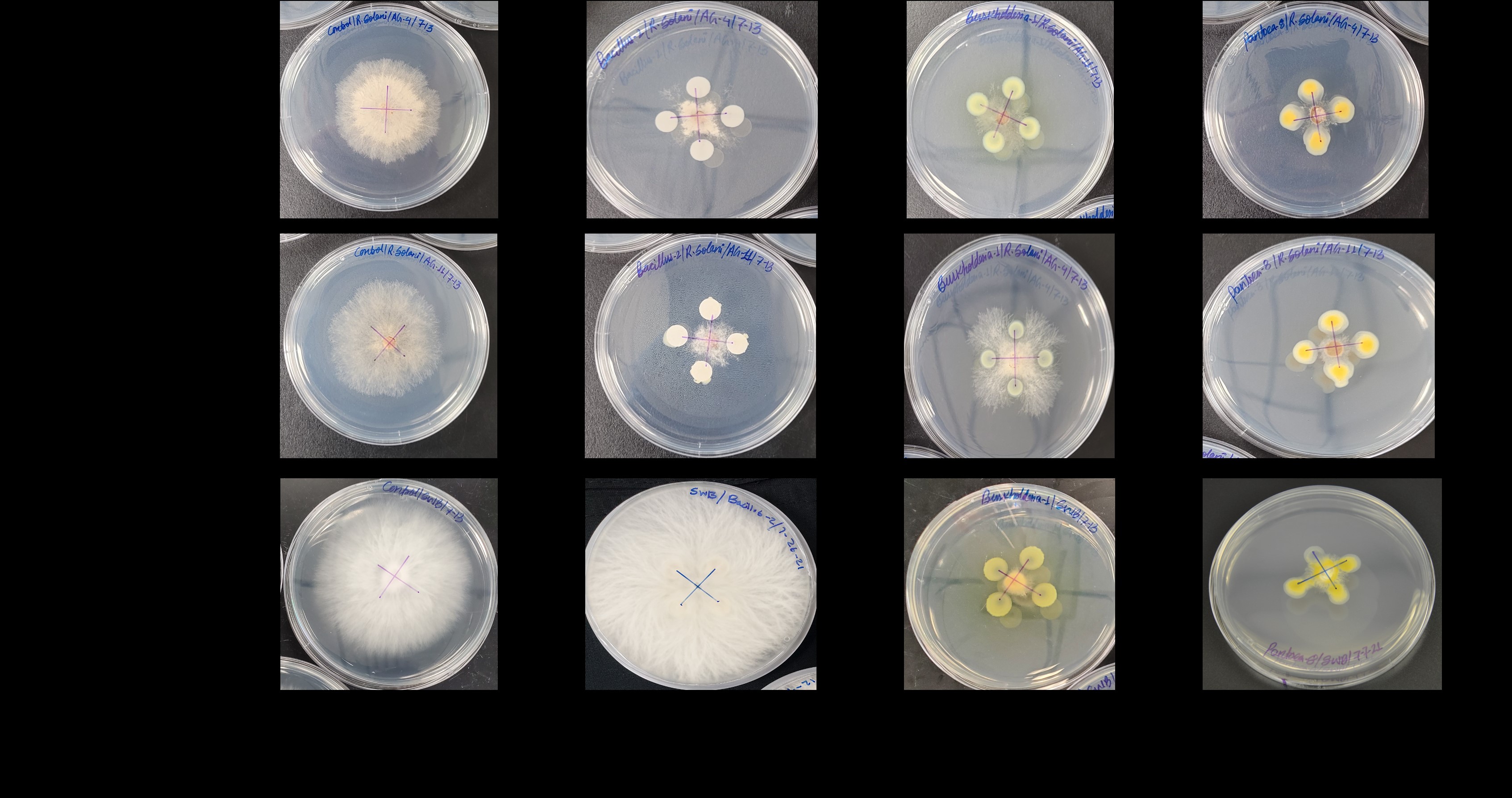

### Supplemental Figure 2

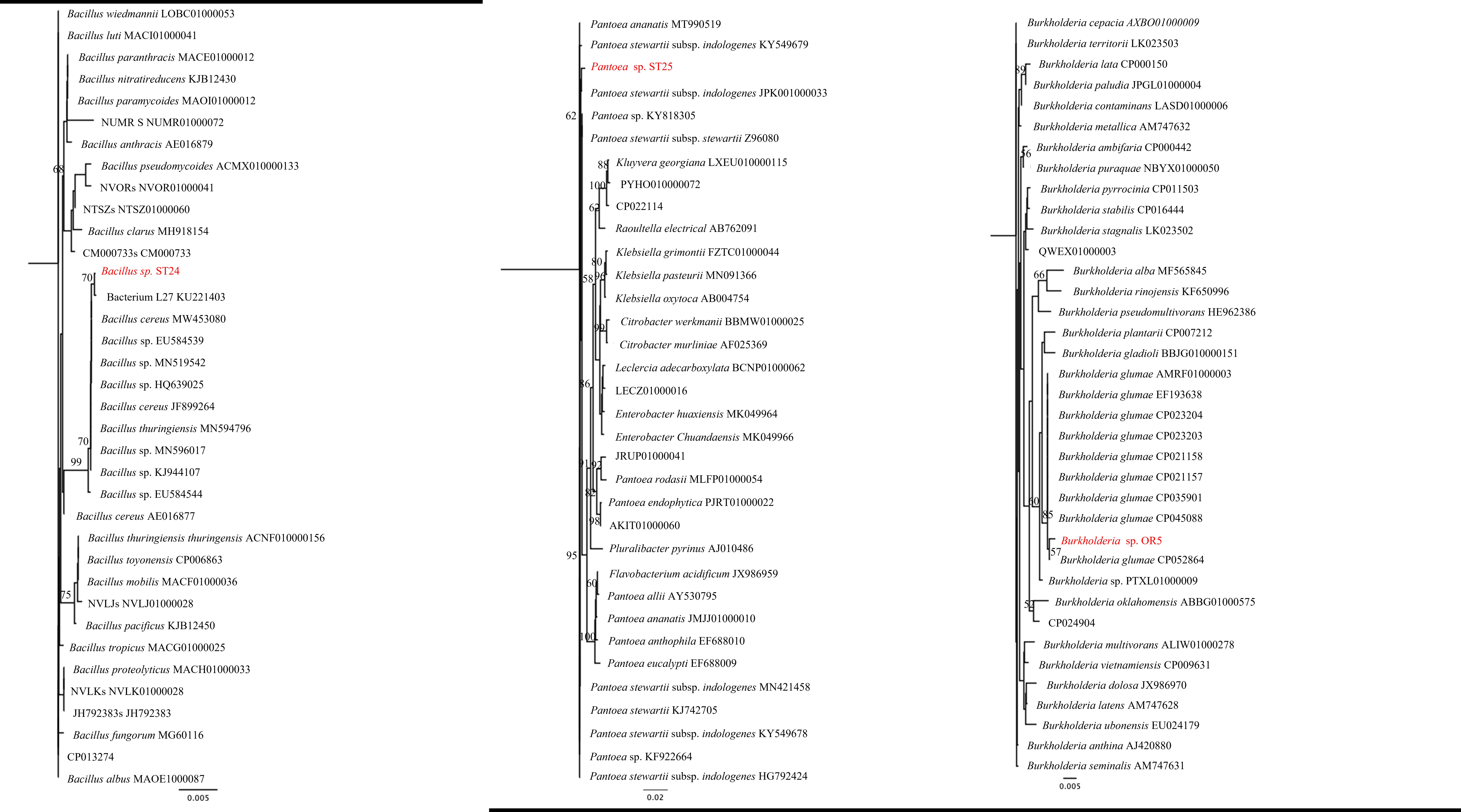
